# The zinc-finger protein POGZ associates with Polycomb repressive complex 1 to regulate bone morphogenetic protein signaling during neuronal differentiation

**DOI:** 10.1101/2025.01.07.631780

**Authors:** Jessenia Chavez, Trevor Wolf, Zhuangzhuang Geng, Yen Teng Tai, Kathleen Bright, James Stafford, Zhonghua Gao

## Abstract

Polycomb Repressive Complex 1 (PRC1) is a family of epigenetic regulators critical for mammalian development. Elucidating PRC1 composition and function across cell types and developmental stages is key to understanding the epigenetic regulation of cell fate determination. In this study, we discovered POGZ, a prominent Autism Spectrum Disorder (ASD) risk factor, as a novel component of PRC1.6, forming the PRC1.6-POGZ complex. Functional assays revealed that POGZ elicits transcriptional repression that is dependent on RING1B expression. Analysis of publicly available ChIP-Seq data showed that POGZ highly colocalizes with RING1B and HP1γ, two PRC1.6 components, at genes involved in multiple aspects of transcriptional regulation in the embryonic mouse cortex. Although *Pogz* knockout (KO) does not compromise stem cell pluripotency, *Pogz* ablation in neuronal progenitor cells (NPCs) led to widespread transcriptomic dysregulation with failed activation of key neuronal genes. Finally, we demonstrated that PRC1.6-POGZ regulates neuronal differentiation by repressing the bone morphogenetic protein (BMP) signaling pathway. These findings reveal a mechanism by which PRC1 and POGZ coordinate transcriptional regulation during neuronal differentiation, which offers insights into how disruptions in this pathway may contribute to neurodevelopmental disorders such as ASD.

## Introduction

Polycomb group (PcG) proteins are key epigenetic modifiers that regulate transcription by modulating chromatin structure and catalyzing histone modifications (1). These functions are essential for critical processes in mammalian development, such as stem cell self-renewal and differentiation (2–6). PcG proteins assemble into two major protein complexes named Polycomb Repressive Complex 1 (PRC1) and 2 (PRC2) (7), which regulate gene expression through distinct repressive chromatin modifications. PRC1 deposits histone H2A mono-ubiquitination (H2AK119ub1), while PRC2 catalyzes H3 lysine 27 mono-, di- and tri-methylation (H3K27me1/2/3) (8, 9).

Mammalian PRC1 complexes display extensive subunit diversity, arising from multiple homologs of core proteins originally identified in *Drosophila melanogaster* (10–12). This diversity includes two E3 ligases (RING1A/B), five chromodomain proteins (CBX2/4/6/7/8), several PH homologs (PHC1-3), and six Polycomb group RING finger proteins (PCGF1-6) (7, 13). PRC1 is further characterized into six major subcomplexes (PRC1.1–1.6), each defined by the exclusive association of one of PCGF1-6 and RING1A/B (14). Additionally, canonical PRC1 complexes associate with CBX and PHC proteins, while non-canonical complexes incorporate RYBP/YAF2 (14–17). Although previous studies laid the groundwork for understanding PRC1 assembly, the distinct roles of subcomplexes in development and differentiation remain relatively unclear.

Specific PRC1 subcomplexes have previously been shown to play unique roles in neurodevelopment and differentiation. For instance, *AUTS2*, a gene often disrupted in individuals suffering from neurological disorders (18, 19), was identified as a component of the PRC1.5-AUTS2 complex in HEK 293 T-REx cells (14). Subsequent studies discovered that PRC1.5-AUTS2 acts as a transcriptional activator in both *in vitro* and *in vivo* models (20–22), thus broadening the repertoire of traditional PRC1 function. Despite these advancements, additional investigation is needed to dissect the possible contributions of other PRC1 subcomplexes in neuronal cells.

*POGO-transposable element with ZNF domain* (*POGZ*) is frequently disrupted in patients with ASD and White Sutton syndrome (23–26). POGZ has been shown to play a role in neurite development in primary cortical neurons (27) and neuronal maturation of iPSC-derived neurons (28). POGZ is also proposed to have transcriptional activity due to its multiple predicted zinc finger domains (29) and sequence-specific DNA-binding properties (30). Indeed, recent research has highlighted a dichotomy in POGZ transcriptional roles, demonstrating its influence on both transcriptional activation and repression (31–34); however, the underlying mechanisms remain unclear.

In this study, we uncovered the surprising interaction between POGZ and PRC1 in primary murine neuronal cells. Biochemical analyses revealed that POGZ specifically associates with the PRC1.6 complex, prompting us to name this complex PRC1.6-POGZ. Functionally, POGZ exhibited transcriptional repression activity that is dependent on RING1B. The formation of the PRC1.6-POGZ complex was further supported by genomic colocalization of POGZ, RING1B, and HP1γ, key components of PRC1.6, in the developing mouse cortex. Interestingly, while *Pogz* is not required for stem cell pluripotency, its loss impairs differentiation to NPCs. Furthermore, during neuronal differentiation, the loss of *Pogz* results in activation of the BMP pathway. These findings uncover an exciting new link between POGZ, a putative ASD risk gene, and PRC1, key epigenetic regulators, in controlling a critical signaling pathway for neurodevelopment.

## Results

### Identification of POGZ as a novel PRC1 subunit in neuronal cells

To explore PRC1 composition in neuronal cells, we employed an unbiased approach using RING1B, the common component of PRC1 complexes, as the bait to identify novel PRC1 interactions. Immunoprecipitation (IP) in nuclear extracts from E17.5 mouse brain, cortical neurons, and neuronal progenitor cells, followed by mass spectrometry, revealed a 50% overlap in the top 50 interactors across the neuronal cell types (**Fig. 1A-B, Table S1**). Among the common interactors were many known PRC1 components, including nearly every PCGF protein (PCGF1/2/3/4/6), multiple CBX proteins (CBX2/4/8), all PHC proteins (PHC1/2/3), RYBP and additional subunits specific to each PRC1.1-1.6 complex (**Fig. 1B**) (14). Particularly interesting was the identification of POGZ, a high-confidence autism risk factor (24–26), as a novel PRC1 interactor (**Fig. 1B**). Prompted by this result, we focused on the association between POGZ and PRC1 to explore how it may influence PRC1 function in neuronal cells. We next sought to determine which of the six PRC1 subcomplexes POGZ associates with. To this end, we performed an IP analysis using our previously generated HEK 293 T-REx cell lines expressing inducible N-terminal FLAG, and HA fused PCGF1-6 (NFH-PCGF1-6) (14). Pull-down with HA beads from each of the PCGF proteins revealed that POGZ specifically associates with the PRC1.6 complex (**Fig. 1C**). A reciprocal FLAG IP using NFH-POGZ inducible HEK 293 T-REx cells confirmed that multiple subunits of PRC1.6, including RING1B, MBLR, HP1γ, and L3MBTL2, co-immunoprecipitated with POGZ (**Fig. 1D**). These data provide evidence for the formation of a novel POGZ-containing PRC1.6 complex, which we name PRC1.6-POGZ.

**Figure 1:**
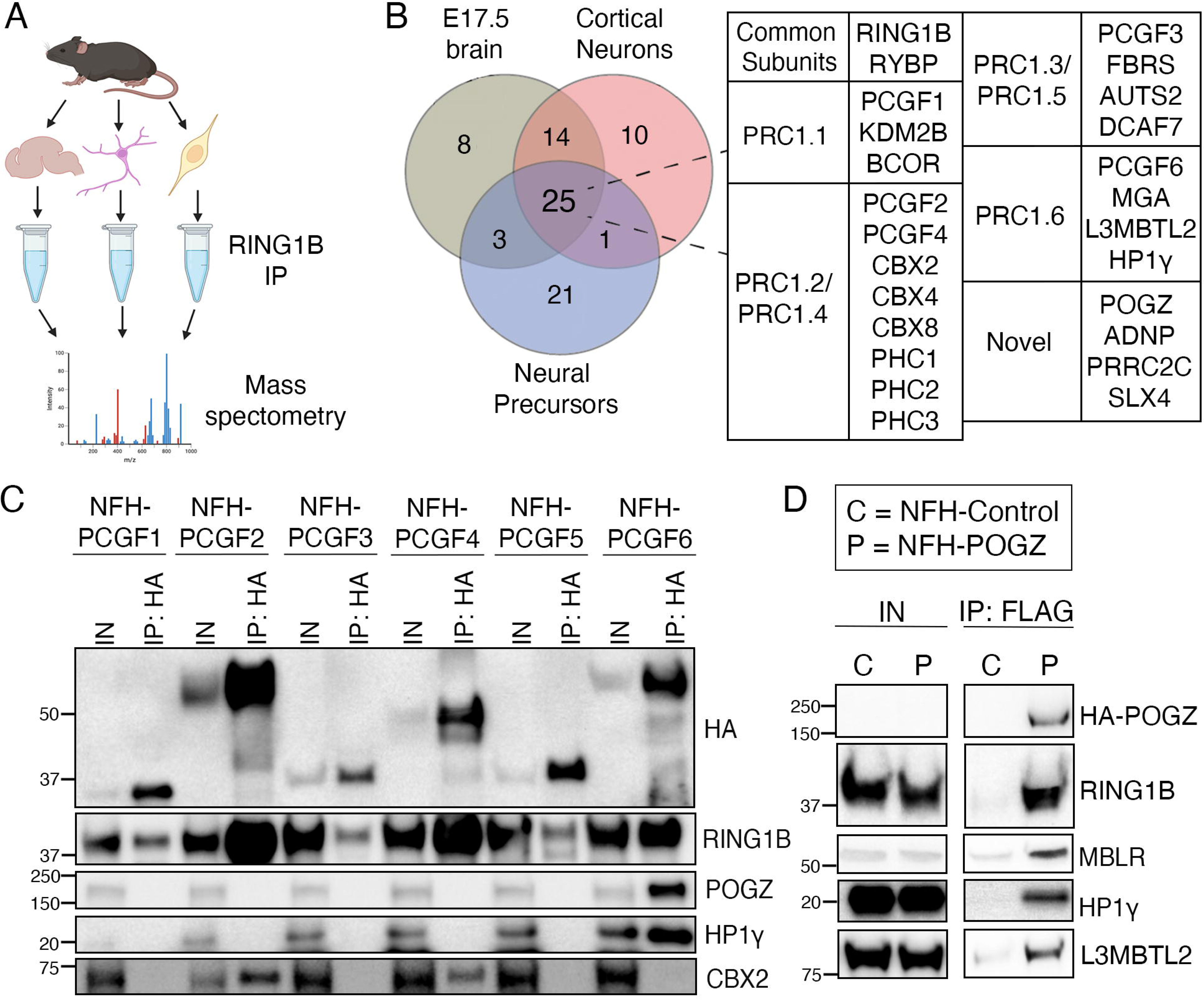
Novel POGZ-Containing PRC1 Complex Identified in the Developing Nervous System. **A.** Schematic displaying the RING1B IP experiment followed by mass spectrometry in (left to right) mouse embryonic brain, cortical neurons, and neuronal progenitors. **B.** RING1B interactors identified in (A) are grouped by their association with specific PRC1 subcomplexes. **C.** HA IP of NFH-PCGF1-6 HEK 293 T-REx cell lines. Bound proteins were resolved on SDS-PAGE and detected by immunoblotting for the indicated antigens. **D.** FLAG IP of NFH-POGZ HEK 293 T-REx cells. The bound proteins were resolved on SDS-PAGE and blotted for indicated PRC1.6 components.

### PRC1.6-POGZ is distinct from other POGZ-containing complexes

To further examine the PRC1.6-POGZ complex, we performed a FLAG-IP using the inducible NFH-PCGF6 cell line followed by glycerol gradient analysis (**Fig. 2A**). Every other fraction was resolved on SDS–PAGE followed by immunoblotting for POGZ and other PRC1.6 components. HA-PCGF6, RING1B, POGZ, HP1γ, and RYBP are enriched in similar fractions, indicating that these factors form a stable complex (**Fig. 2B**). Previous reports have suggested that POGZ associates with both the ChAHP complex, a recently identified transcriptional repressor complex required for embryonic stem cell (ESC) maintenance (35), and the esBAF complex, a transcriptional activator complex in ESCs that maintain stem cell identity (30). To explore POGZ association with epigenetic complexes other than PRC1, we performed glycerol gradient analysis using NFH-HP1γ, the shared subunit among the PRC1.6-POGZ, ChAHP, and esBAF complexes. Our results revealed that HA-HP1γ, RING1B, POGZ, and additional PRC1.6 specific components, L3MBTL2 and WDR5, were enriched in the same fractions, further supporting the formation of the novel PRC1.6-POGZ complex (**Fig. 2C**). In contrast, CHD4, a core subunit of the ChAHP complex, and BRG1, the core ATPase subunit of the esBAF complex, were enriched in earlier fractions compared to those corresponding to the PRC1.6-POGZ complex (**Fig. 2C**). Taken together, our data strongly suggests the existence of a new POGZ-associated complex that constitutes a subtype of PRC1 (**Fig. 2D**).

**Figure 2.**
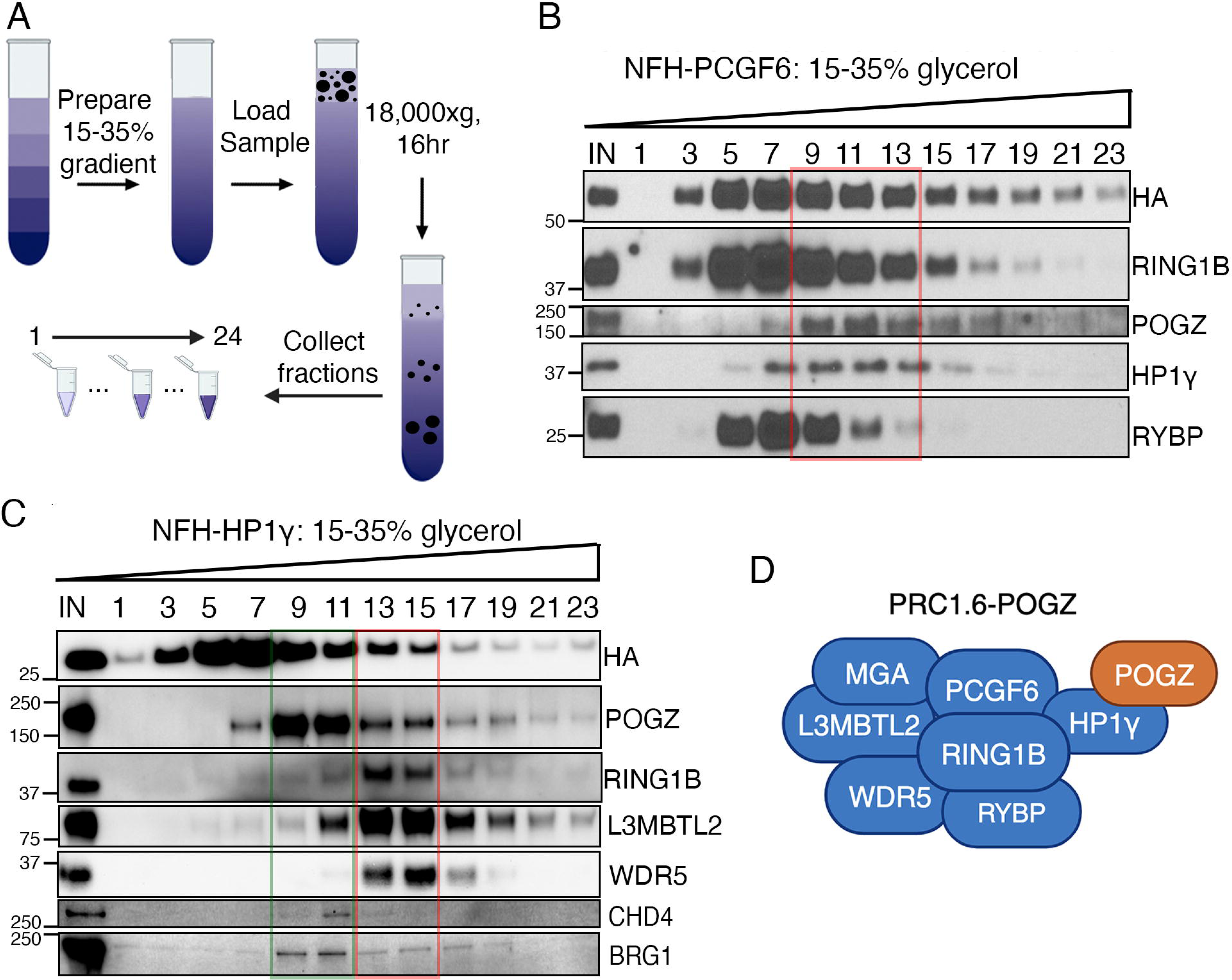
POGZ is a stable component of the PRC1.6 complex. **A.** Schematic of the glycerol gradient experimental workflow. **B-C.** Glycerol gradient analysis of FLAG-purified PCGF6 (B) and HP1γ (C) complexes. Every other fraction from a 15–35% glycerol gradient was resolved on SDS-PAGE, followed by immunoblotting for the indicated antigens. Red boxes highlight fractions containing PRC1.6-POGZ subunits, and the green box corresponds to non-PRC1 POGZ-containing complexes. **D.** Schematic illustration of the proposed PRC1.6-POGZ complex. Blue subunits represent previously identified PRC1.6 subunits, and the orange subunit highlights POGZ as a novel PRC1.6 interactor.

### POGZ-Dependent Repression Requires RING1B

PRC1.6 functions as a transcriptional repressor (36). To investigate the role of PRC1.6-POGZ in transcriptional regulation, we performed luciferase assays using a stable HEK 293 T-REx cell line containing an integrated luciferase reporter with five consecutive GAL4 DNA binding sites (UAS) and doxycycline-inducible GAL4–POGZ (**Fig. 3A**). We also used previously established lines with GAL4 alone or GAL4-PCGF4 (20) as the negative and positive controls, respectively. Upon doxycycline induction, luciferase activity decreased in GAL4–PCGF4 cells (**Fig. 3B**), aligning with its function in transcriptional repression (20). Consistent with previous reports, induced expression of GAL4-POGZ also led to a significant decrease in luciferase activity (**Fig. 3B**), supporting its role as a transcriptional repressor (33). To further investigate whether POGZ association with PRC1 was essential for its transcriptional repression activity, we silenced RING1B through short interfering RNAs (siRNAs) in GAL4–POGZ cells. Interestingly, RING1B knockdown (KD) in GAL4–POGZ cells significantly reduced repression of reporter gene transcription (**Fig. 3C-D**). These findings demonstrate that the transcriptional repressor function of POGZ requires its association with RING1B, highlighting a key mechanistic link between POGZ and PRC1 in transcriptional repression.

**Figure 3.**
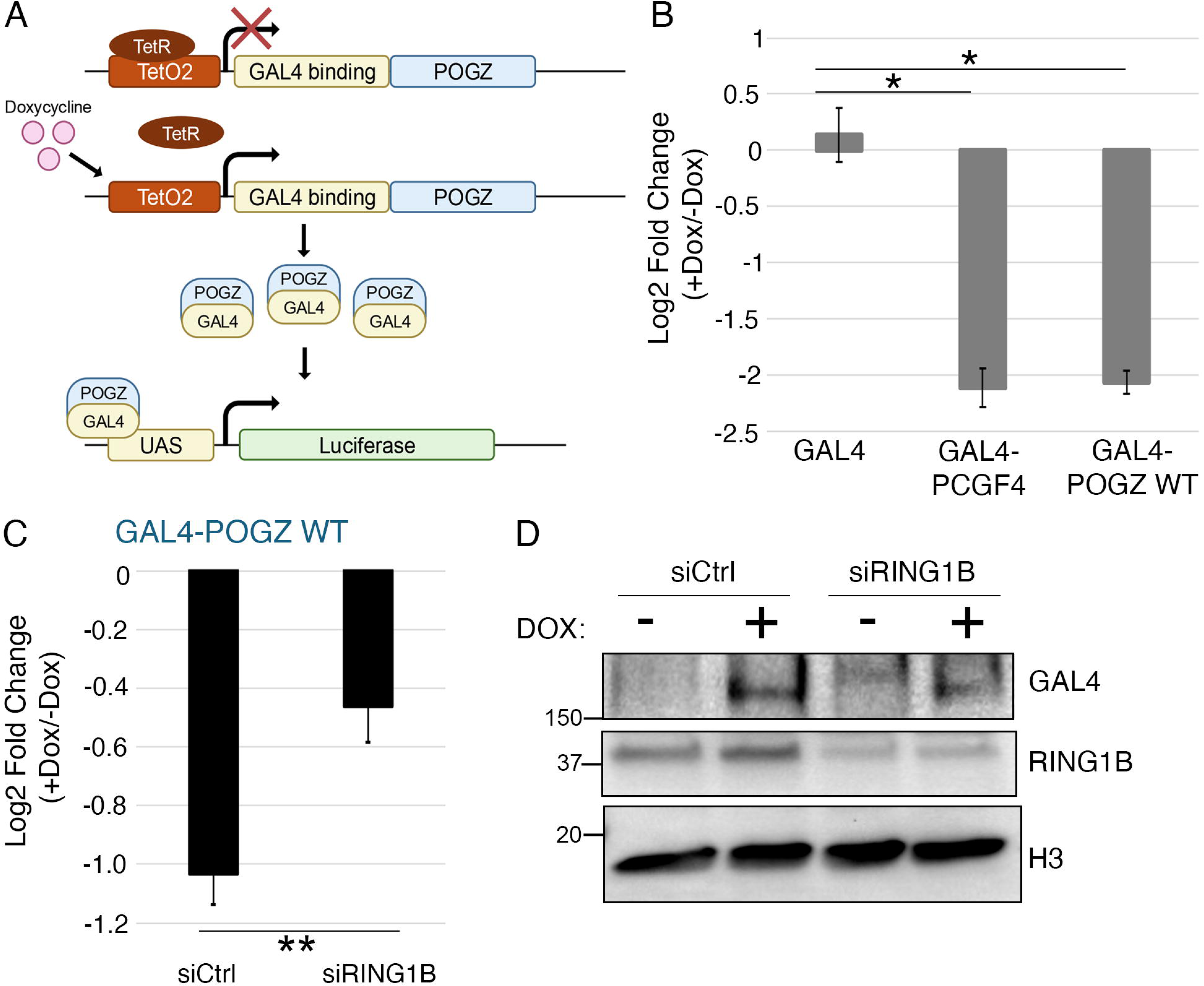
RING1B Activity Is Essential for POGZ-Mediated Transcriptional Repression. **A.** Schematic of the luciferase reporter system used to examine the transcriptional impact of POGZ. **B.** Fold change in luciferase activity in cells expressing GAL4 only, GAL4–PCGF4, and GAL4–POGZ after 24-hour doxycycline induction. Each value is the mean of three independent measurements, with error bars representing the standard deviation. \**p*<0.05, \*\**p*<0.01 by two-sided t-test, compared with the GAL4 control. **C.** Fold change in luciferase activity in GAL4–POGZ cells upon KD of RING1B. Cells were transfected with control or RING1B siRNAs, and luciferase activity was measured after 24-hour doxycycline induction. Each value is the mean of three independent measurements, with error bars representing the standard deviation. \**p*<0.05, \*\**p*<0.01 by two-sided t-test, compared with the GAL4 control. **D.** Immunoblotting of samples used for luciferase activity reporter assay in (C) to demonstrate NG4-POGZ WT induction and RING1B KD efficiency.

### POGZ colocalizes with PRC1.6 at neurodevelopmental genes in mouse cortex

With the identification of POGZ as a PRC1.6-associated subunit, we next sought to investigate whether POGZ colocalizes with PRC1.6 in a developmentally relevant model. To address this, we utilized existing ChIP-seq datasets of POGZ, RING1B, and HP1γ in mouse cortex (31, 37). Our analysis identified 2,927 total POGZ peaks, with 46.3% of peaks (n = 1358) overlapping with both RING1B and HP1γ, therefore representing PRC1.6-POGZ target loci (**Fig. 4A**). We also found that 45.9% of POGZ peaks (n = 1344) overlapped with only HP1γ, possibly representing loci targeted by non-PRC1.6-POGZ-containing complexes (**Fig. 4A**). Only a small percentage of peaks are co-occupied by both POGZ and RING1B (1.7%, n=51) or POGZ alone (6%, n=174) (**Fig. 4A**). Gene ontology (GO) analysis of PRC1.6-POGZ targets genes revealed enrichment for terms relating to transcriptional regulation (**Fig. 4B**). We observed strong POGZ, RING1B, and HP1γ localization at *Klf6* and *Myh9*, which are PRC1.6-POGZ target genes that are implicated in various aspects in neurodevelopment (**Figure 4C**) (38–41). These findings highlight the complexity of POGZ genomic occupancy and identify the genes that are targeted by the PRC1.6-POGZ complex.

**Figure 4.**
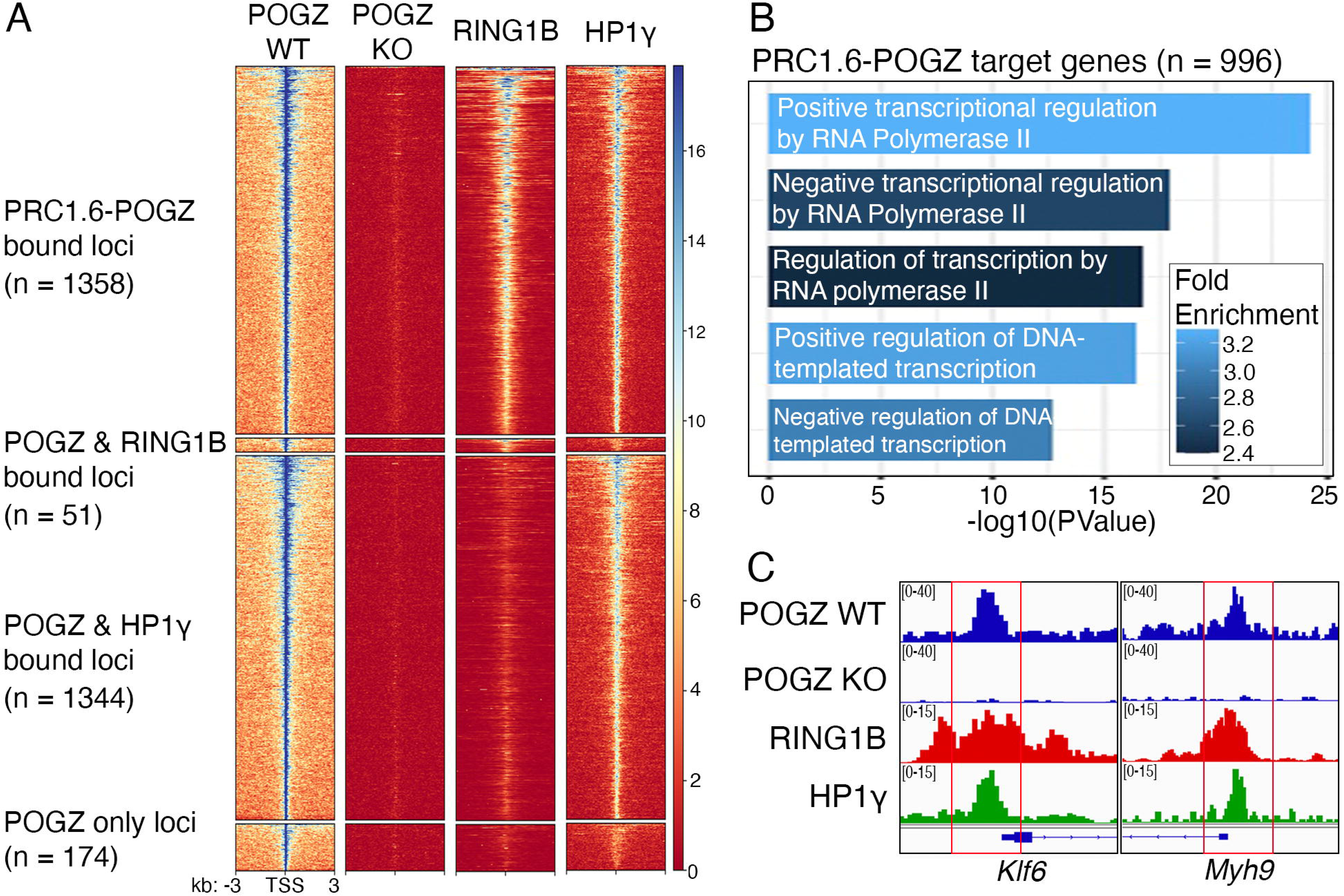
PRC1.6-POGZ genomic localization within embryonic mouse cortex. **A.** Heatmap showing enrichment of PRC1.6-POGZ, POGZ & RING1B, POGZ & HP1γ, and POGZ only bound peaks centered around TSS (±3 kb) regions across loci. **B.** GO analysis of PRC1.6-POGZ target genes in mouse cortex. The x-axis (in logarithmic scale) corresponds to the binomial raw P values. **C.** Representative genomic tracks showing the normalized tracks of POGZ, RING1B, and HP1γ at the indicated loci.

### *Pogz* deletion leads to a transcriptomic alteration in NPCs

To investigate the impact of PRC1.6-POGZ on its target gene expression, we generated a *Pogz* KO ESC line (**Fig. S1A**). Deletion of *Pogz* was confirmed by Sanger sequencing, immunoblotting, and RT-qPCR (**Fig. S1A-C**). Examination of pluripotency markers *Nanog* and *Oct4* at both protein and transcript levels showed no changes in expression following *Pogz* KO (**Fig. S1B-C)**. Additionally, alkaline phosphatase (AP) staining of *Pogz* WT and KO ESCs revealed no noticeable difference in AP activity, further indicating that *Pogz* deletion does not affect pluripotency (**Fig. S1D**).

Using an established differentiation protocol (42, 43), *Pogz* WT and KO ESCs were differentiated into NPCs (**Fig. 5A**). To evaluate the impact of *Pogz* KO on neuronal differentiation, we examined the expression of NPC marker genes, including *Pax6*, *Nes*, and *Sox1,* using RT-qPCR. We found that these marker genes failed to be induced in *Pogz* KO NPCs, pointing to a disruption in the differentiation process (**Fig. 5B**). To further examine the changes in gene expression upon loss of *Pogz,* we performed RNA-Seq in *Pogz* WT and KO ESCs and NPCs (**Fig. 5C**). Our RNA-Seq analysis identified many differentially expressed genes between *Pogz* WT and KO NPCs, with 1,139 downregulated and 1,973 upregulated genes identified in *Pogz* KO NPCs (**Fig. 5C-D**). GO analysis showed that these downregulated genes were enriched in terms related to transcriptional regulation, differentiation, and neurogenesis (**Fig. 5D**). GO analysis of the up-regulated genes showed enrichment in terms related to general metabolism, macromolecule transport, and angiogenesis (**Fig. 5D**). These results demonstrate that *Pogz* is required for ESC neuronal differentiation through establishing an NPC specific transcriptome.

**Figure 5.**
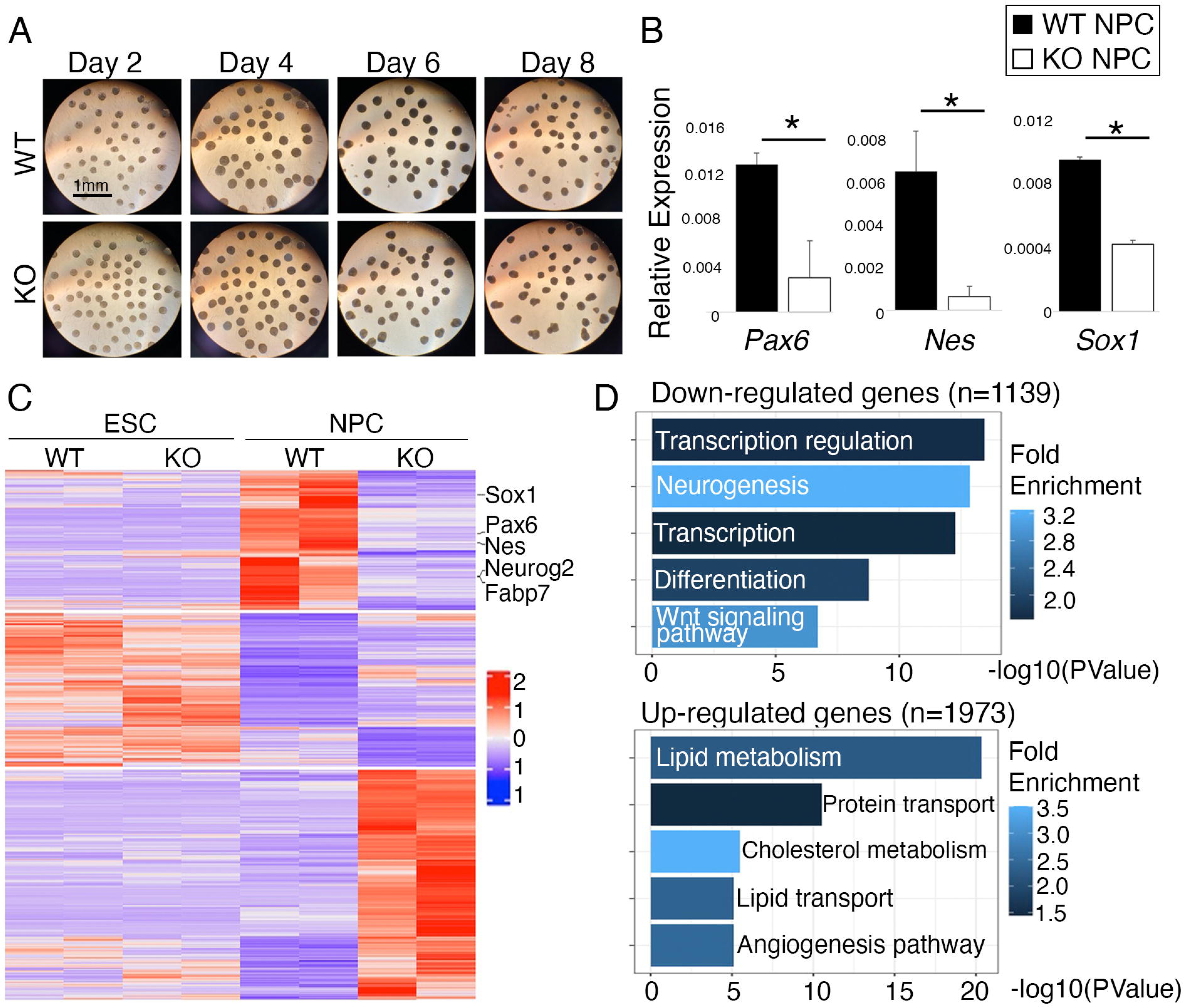
*Pogz KO l*eads to dysregulated NPC transcriptome. **A.** Brightfield images of *Pogz* WT and KO cells at various time points of the differentiation process. **B.** RT-qPCR analysis of NPC marker gene expression in *Pogz* WT and KO ESCs and NPCs. All mean values and standard deviations were calculated from three independent measurements. * *p*<0.05, ** *p*<0.01, *** *p*<0.001. **C.** Heatmap of the global transcriptomic profile in *Pogz* WT and KO ESCs and NPCs. Duplicate samples were subjected to RNA-seq analysis. TPM values for each gene were converted to z-scores and used to generate the heatmap. **D.** GO analysis of down-regulated (top) and up-regulated (bottom) genes in *Pogz* KO NPCs compared to *Pogz* WT NPCs. The x-axis (in logarithmic scale) corresponds to the binomial raw P values.

### PRC1.6-POGZ negatively regulates the BMP pathway during neuronal differentiation

To further understand the impact of *Pogz* KO on gene regulation during neuronal differentiation, we cross-referenced ChIP-Seq data from mouse cortex with our RNA-Seq results. Our analysis revealed 68 up-regulated and 60 down-regulated PRC1.6-POGZ target genes (**Fig. 6A, Table S2-S3**). Considering that POGZ is a known transcriptional repressor (33, 34) and its repressive activity is dependent on RING1B (**Fig. 3**), we focused on PRC1.6-POGZ target genes that were de-repressed in *Pogz* KO NPCs. Among the de-repressed genes, our RNA-Seq data and RT-qPCR validation revealed that *Klf6*, a Transforming Growth Factor beta (TGF-β) activator, expression was approximately three-fold higher (fold-change = 3.1) in *Pogz* KO NPCs than in WT NPCs (**Fig. 6A-B**). During neuroectoderm differentiation, the TGF-β and BMP signaling pathways must be inhibited as they promote meso- and endodermal differentiation (44–46). The activation of the TGF-β/BMP pathway leads to the phosphorylation of regulatory SMAD (R-SMAD) proteins, among which SMAD1/5/9 respond to BMP and SMAD2/3 respond to TGF-β (44).

**Figure 6.**
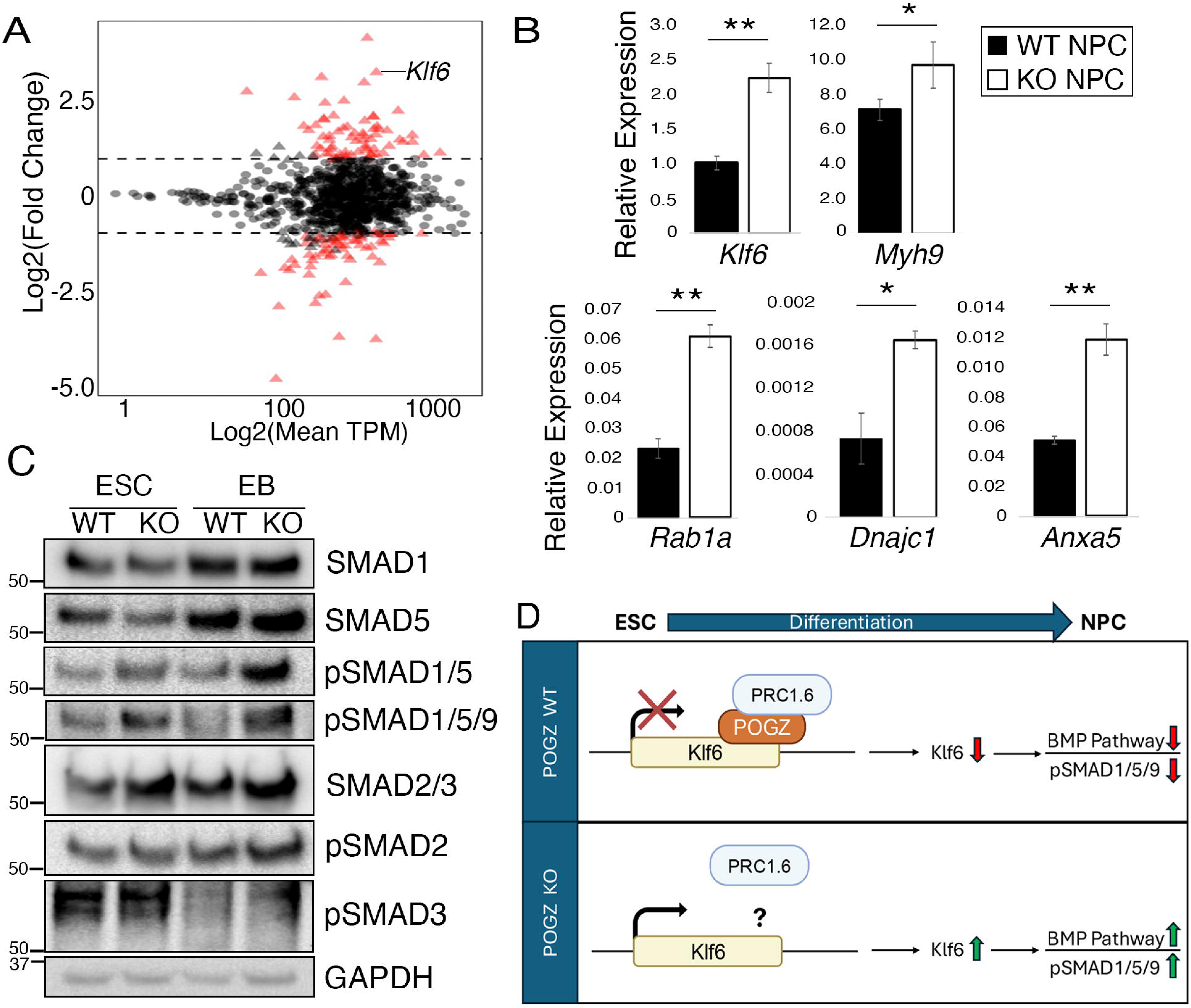
*Pogz* KO leads to hyperactivation of BMP signaling. **A.** Comparison of PRC1.6-POGZ target gene expression levels based on RNA-seq analysis from *Pogz* WT and KO NPCs. The x-axis is the log_2_ value of the average TPM value of a gene, and the y-axis is the log_2_ value of fold changes of TPM of a gene between two groups. Triangles and circles represent data points with absolute log_2_ fold change greater or less than 1, respectively. Pink data points highlight genes with an absolute log_2_ fold change > 1 and an adjusted p-value <0.05. Grey indicates genes that do not meet both thresholds. **B.** RT-qPCR analysis of PRC1.6-POGZ target gene expression in *Pogz* WT and KO NPCs. **C.** Immunoblotting of *Pogz* WT and KO ESC and EB using the indicated antibodies. **D.** Model for the role of PRC1.6-POGZ regulation of the BMP pathway during neuronal differentiation.

The observed upregulation of *Klf6* expression in *Pogz* KO NPCs led us to hypothesize that increased expression of *Klf6* alters TGF-β/BMP signaling pathways in these cells. Our immunoblotting experiments revealed that loss of *Pogz* led to an increase in phosphorylated SMAD1/5/9 (pSMAD1/5/9) at the ESC and EB stages compared to WT cells (**Fig. 6C**). Conversely, the TGF-β pathway does not play a significant role in the context of neuronal differentiation of *Pogz* KO cells as there was little to no change in pSMAD2/3 levels in KO cells (**Fig. 6C**). These findings suggest that *Pogz* is required for inhibiting BMP signaling through PRC1.6-POGZ-mediated repression of *Klf6*, which in turn contributes to the regulation of proper neuronal differentiation.

## Discussion

Our investigation into the composition of PRC1 in neuronal cells led to the discovery of a stable PRC1.6-POGZ complex consisting of a novel component, POGZ, an autism risk factor (24–26). The existence of the PRC1.6-POGZ complex is further supported by genomic colocalization of POGZ, RING1B, and HP1γ in the mouse cortex. We further demonstrated that RING1B mediates POGZ transcriptional repression and that *Pogz* KO NPCs exhibit de-repression of its target genes. Our results suggest a model where PRC1.6-POGZ controls proper neuronal differentiation by inhibiting BMP signaling (**Fig. 6D**).

Our discovery of POGZ as a novel interactor of PRC1.6 in neuronal cells is particularly intriguing, given the well-established role of PCGF6 as a master regulator of ESC pluripotency (36, 47–49). This finding raises the possibility of alternative PRC1.6 complex composition in differentiated cell types. Previous studies have identified various DNA-binding factors as PRC1 recruiters in a cell-type-dependent manner (50–53). PRC1.6 is known to be recruited to target loci through its interactions with a variety of chromatin-associated factors, including E2F6/DP1 and MGA/MAX (47, 54–57). It is worth pointing out that although we identified MGA as a RING1B interacting protein in neuronal cells, E2F6, DP1, and MAX were not present in our MS analysis (**Fig. 1B**). It is highly likely that POGZ serves as a PRC1.6 recruiter to target genes in neuronal cells by leveraging its zinc finger domains (29) and sequence-specific DNA-binding properties (30) to mediate recruitment. Future studies, including loss-of-function experiments, will be needed to fully dissect the dynamics of PRC1.6-POGZ recruitment in neuronal cells.

TGF-β signaling plays an important role in cell fate determination (44). The differentiation of ESCs into neuroectodermal lineage requires the absence of TGF-β and its family members, BMPs, while the mesoderm and endoderm differentiation need this signaling (45). Furthermore, inhibition of TGF-β and BMP signaling greatly enhances the efficiency of neuronal differentiation from ESCs (46). Interestingly, previous studies on PRC1 and related complexes revealed their critical role in regulating these signaling during neuronal differentiation (58, 59). Our study led to the discovery of the role of PRC1.6-POGZ in repressing BMP signaling by targeting *Klf6* (**Fig. 4 and 6**), a previously reported activator for the TGF-β pathway (Kojima et al., 2000). Although the exact mechanism by which PRC1.6-POGz represses *Klf6* remains to be fully understood, our model suggests that in WT cells undergoing neuronal differentiation, POGZ recruits PRC1.6 to the *Klf6* loci, leading to its repression and inhibition of the BMP pathway (**Fig. 6D**). These studies clearly establish a link between various PRC1 complexes and ASD risk factors and their impact on the deregulation of TGF-β/BMP pathways as a potential shared mechanism in neurological disorders. These findings provide a promising avenue for the development of targeted therapies.

## Experimental procedures

### Isolation of neuronal progenitors and cortical neurons

Neuronal progenitors and cortical neurons were isolated as previously described (60, 61). Briefly, to obtain neuronal progenitors, the mouse brains were removed and sliced into 1.0 mm coronal sections. Slices containing the hippocampus and the SVZ were placed into a 10cm petri dish filled with ice-cold dissection buffer (Hank’s balanced salt solution, 2 mM HEPES). Using a microscope, the dentate gyrus and SVZ were isolated from relevant slices, minced in dissection buffer, and placed in a 15ml tube. To obtain cortical neurons, the cortex was isolated from mouse brains, minced in the dissection buffer, and placed in a 15ml tube. The following procedure was used to isolate neuronal progenitors and cortical neurons in their respective tissue samples. After mincing, tissue samples were spun down at 200g for 1min at RT. In a cell culture hood, dissection buffer was aspirated, and 1mL of 0.05% Trypsin-EDTA was added to the tissue samples and incubated at 37°C with slow rotation for 15-20 minutes, followed by gentle resuspension. Wash media (DMEM, B27 without vitamin A, GlutaMAX, and Antibiotic-Antimycotic) was added to stop digestion, and the samples were incubated at 37°C with slow rotation for 15-20 minutes, followed by additional resuspension to break tissue chunks into single cells. Cells were washed twice by adding 8 mL of wash media followed by centrifugation at 200 g for 5 min. Cells were resuspended in 1mL of growth media and plated into a single well of a low-adherent 24-well culture plate. The cells were incubated in a 5% CO_2_ 37°C incubator for 48 hr. After the initial 48 hours, half of the media was replaced with fresh growth media every second day. Cells were collected after 1-2 weeks of culturing.

### RING1B IP and Mass Spectrometry

To conjugate antibodies to beads, a 3:1 Protein A/G mixture was washed using 1X PBS. Beads were then incubated with 100 µg of antibody for 3 hours at room temperature (RT) or overnight (O/N) at 4°C. The antibodies used are RING1B (Bethyl, cat. no. A302-869A-T) and IgG (Cell Signaling, cat. no. 2729S). Beads were then washed twice with 0.2M Sodium borate buffer, pH 8.0. Crosslinking was performed using 20 mM DMP (made in 0.2M sodium borate buffer) followed by rotation for 2 hours at RT. Beads were centrifuged and washed three times with 0.2M ethanolamine-HCl, pH 8.0, to stop the reaction. Next, 0.1M glycine, pH 2.5, was added to beads, followed by washes using 1M Tris, pH 8.0. Beads were washed three times with PBS.

To perform the IP, E17.5 brain, cortical neurons, and neuronal progenitor cells were extracted from mice and processed using the nuclear extraction protocol below. 1mg of nuclear extract and beads were incubated O/N at 4°C with rotation. The lysate-bead mix was then centrifuged and washed with Buffer C three times with rotation at 4°C. For the last wash, the beads were resuspended in 1X PBS and transferred to a Micro-spin column (Pierce, cat no. 89879). Samples were centrifuged, then eluted with 0.1M glycine, pH 2.5, and neutralized with 10µL of 1M Tris, pH 9.5. Each elution is performed with an equivalent volume of glycine to beads. Elution was repeated three times. Elutions were run using the SilverQuest™ Silver Staining Kit according to manufacturer instructions (Invitrogen, cat. no. LC6070), and samples were submitted for semi-quantitative mass spectrometry.

### Nuclear extraction

The IP and glycerol gradient experiments described herein were performed using nuclear extracts (NE). NEs were prepared as previously described (14). Briefly, the buffers used for nuclear extraction are Buffer A (10mM Tris-HCl, pH 7.9, 1.5 mM MgCl_2_, 10 mM KCl) and Buffer C (20mM Tris-HCl, pH 7.9, 1.5 mM MgCl_2_, 420mM NaCl, 0.2mM EDTA, 25% glycerol). Protease inhibitors were added before every experiment at the following concentrations: 0.2 mM PMSF, 1 µg/ml Pepstatin A, 1 µg/ml Leupeptin, and 1 µg/ml Aprotinin. Cell pellets were first resuspended using Buffer A, incubated on ice for 15 minutes, and homogenized using a 2mL dounce homogenizer for 15 strokes. After 30min rotation at 4°C, the samples are centrifuged at 15,000 rpm for 15 min at 4°C. The supernatant (cytosolic fraction) was removed, and the remaining pellet was resuspended in Buffer C. After resuspension, the lysate was homogenized, rotated at 4°C for 30 min, and centrifuged at 15,000rpm for 15min. The resulting supernatant consisted of the NE and was then subjected to various downstream analyses.

### Cell lines

HEK 293 T-REx cells stably expressing pINTO-NFH-tagged proteins were generated and maintained as previously described (14). Briefly, pINTO-N-FH-fusion proteins or vector control (pINTO-NFH) were transfected into HEK 293 T-REx cells using Lipofectamine 3000 (Invitrogen) according to manufacturer’s protocol. Cells underwent selection using 200 µg/mL zeocin and 10 µg/mL blasticidin. After selection, cells were maintained in medium containing 50-100 µg/mL zeocin and 10 µg/mL blasticidin. Luciferase reporter cell lines expressing GAL4-fusion proteins or vector control (pINTO-GAL4) were generated and maintained as previously described (14). Briefly, GAL4-fusion proteins or vector control (pINTO-GAL4) were transfected into HEK 293 T-REx -luciferase cells containing a stably integrated 5XGal4RE-tk-Luc-neo construct and selected with 200 µg/ml zeocin, 10 µg/ml blasticidin, and 400 µg/ml G418 (62). After selection, cells were maintained in medium containing 50-100 µg/mL zeocin and 10 µg/mL blasticidin. All HEK 293 T-REx cell lines were maintained in standard DMEM medium containing 10% FBS (Atlanta Biologicals, Cat# S11050), L-glutamine, and penicillin/streptomycin.

### Immunoprecipitation

Immunoprecipitation experiments were performed as described previously (14). To perform immunoprecipitation, NEs are incubated with pre-washed FLAG M2 beads (Sigma, cat. no. A2220) O/N at 4°C. After the overnight rotation, the beads were washed five times with Buffer W (1/3 volume of Buffer A, 2/3 volume of Buffer C, 0.02% IGEPAL, 0.2 mM PMSF, 1 µg/ml Pepstatin A, 1 µg/ml Leupeptin and 1 µg/ml Aprotinin). To elute, 40µL of BC and 10µL of 5X SDS were added to each sample and boiled for 5min at 95°C. Samples were centrifuged, and the supernatant was collected for immunoblotting.

### Glycerol gradient analysis

Glycerol gradient analysis was performed as previously described (14). To perform glycerol gradient analysis, 30-15cm plates per HEK 293 T-REx cell line were treated with 100µg/mL doxycycline for 24 hours. The resulting pellet was processed to obtain the NE (see above). Volumes for BA and BC were adjusted to 15ml and 13.5ml, respectively. 12 ml NE was mixed with 3 ml Buffer A, 0.02% NP-40, and 200µL of pre-washed FLAG M2 beads. After rotation at 4°C O/N, the M2 beads were washed five times with Buffer W and then eluted with 500µL of 250 µg/ml FLAG peptides in Buffer W by rotating at 4°C for 1 hour. The M2 eluate was added on the top of a 4.5 ml 15-35% glycerol gradient and centrifuged in an SW60Ti rotor (Beckman) at 18,000xg at 4°C for 16 hours. The resulting gradient was then fractionated every 180 µl. All odd-numbered fractions were then subjected to SDS-PAGE and immunoblotting.

### Luciferase reporter assay

To perform the luciferase assays, cell lines were plated in 6-well plates in sextuplicate at 300,000 cells per well on Day 0. Three of the wells were treated with either 100µg/mL doxycycline or vehicle on Day 1 for 24 hours. The cells were then assayed on Day 2 using Promega’s Luciferase Assay System (cat. no. E1500) per manufacturer’s instructions, with the exception that the luciferase assay reagent was added manually. Luminescence was measured using the Promega GloMax 96 Microplate Luminometer (cat. no. E6501). Luciferase results were normalized to protein concentration and then analyzed by comparing luminescence fold change of induced vs. non-induced conditions on a per-cell line basis. Fold changes were then compared by a two-sample t-test.

### RNA interference

Cells were transfected with siRNAs using Lipofectamine RNAiMAX (Life Technologies) according to the manufacturer’s protocol. Human siRNAs used in this study are si-ctrl (Qiagen, AllStars Negative Control siRNA, cat no. SI03650318) and si-RING1B (Qiagen, cat no. SI00095543).

### ChIP-Seq analysis

Published ChIP-seq data for POGZ in wild-type and knock-out cortex and HP1γ in wild-type cortex were obtained from NCBI GEO Series GSE187010 (31). Published ChIP-seq data for RING1B in wild-type cortex were also obtained from the DNA Data Bank of Japan Sequence Read Archive (accession no. DRA010296) (37). Sequencing results were mapped to the mm10 using BWA (63). Duplicated reads were removed with Samtools (64). BamCoverage and computeMatrix from deepTools were used to generate the normalized Bigwig files and ChIP-Seq heatmaps, respectively (65). Macs2 callpeak was used to generate narrowPeak files (66). Bedtools Intersection was used to identify PRC1.6-POGZ, POGZ & RING1B, POGZ & HP1γ, and POGZ-only bound loci (67). ChIP-seq read density files were generated using igvtools and were viewed in Integrative Genomics Viewer (IGV) (68). Gene annotations were obtained with genomic regions enrichment of annotations tool (GREAT) (69). GO analysis was performed using DAVID (70).

### CRISPR/Cas9-mediated gene editing

Oligos corresponding to candidate sgRNA sequences were purchased from IDT and cloned into pX458 (Addgene) as previously described (71). The sgRNA sequences used were: sgRNA1 (5’-ACTTGTGGGACGCCAACTGT<u>CGG</u>*-*3’) and sgRNA2 (3’-CTAGTTTGGGATTCGAGGTC<u>TGG</u>-5) (PAM trinucleotides are underlined). Plasmids carrying sgRNAs were transfected into ESCs using Lipofectamine (Life Technologies). Two days after transfection, cells were sorted into 96-well plates coated with gelatin at one cell per well. Clones were tested by genomic PCR using primers: 5′-TTCCTTTTGTTCAGTGGACAGC-3′, and 3′-AAGGAGGGAGCTACATTGACC-5′. Positive clones were further validated by Sanger sequencing and immunoblotting using a POGZ antibody (Bethyl, cat# A302-510A).

### ESC culturing

ESCs and CRISPR-engineered ESCs were cultured in embryonic stem cell medium comprised of DMEM medium, supplemented with 15% FBS (ES certified, Atlanta Biologicals, cat. no. S10250), LIF, non-essential amino acids, β-mercaptoethanol, L-glutamine, penicillin/streptomycin, sodium pyruvate, and two small-molecule kinases (MEK and GSK3) inhibitors (PD0325901, Cayman Chemical, cat. no. 13034 and CHIR99021, Cayman Chemical, cat. no. 13122). When reviving from liquid nitrogen, ESCs are initially plated on mitotically arrested mouse embryonic fibroblast feeder cells. ESCs are split every 48 hours on gelatin-coated plates.

### Neuronal differentiation

Neuronal differentiation was performed as previously described (42, 43). ESCs cultured on mouse embryonic fibroblasts were split into gelatin-coated plates for two passages before differentiation. The differentiation medium contained DMEM medium, 15% FBS, non-essential amino acids, β-mercaptoethanol, L-glutamine, penicillin/streptomycin, and sodium pyruvate. On Day 0, ESCs differentiated using the hang-drop method. Each 20uL droplet contained ∼1500 ESCs. The hang drop plates were placed in a 37°C incubator with 5% CO_2_ content. On day 2, the ESCs will be differentiated into EBs. EBs were then transferred to a suspension plate containing differentiation medium for two additional days. On Day 4, 5µM retinoic acid was added to the culture medium to trigger differentiation towards the neuronal lineage. By Day 8, NPCs will have been generated. NPCs were then washed with 1X PBS, pelleted, and collected for various downstream analyses.

### RNA-Seq and analysis

cDNA libraries were prepared using the NEXTflex™ Illumina Rapid Directional RNA-Seq Library Prep Kit (Illumina, cat. no. NOVA-5198-0) according to the manufacturer’s instructions. Libraries were loaded onto a TruSeq Rapid flow cell on an Illumina HiSeq 2500 (Genome Sciences Facility, Penn State College of Medicine). The samples were run for 50 cycles using either a single-read or pair-end recipe according to the manufacturer’s instructions. The gene expression abundance was quantified using Kallisto against mm10 transcriptome reference (72). Differentially expressed genes (DEG) analysis was identified using DESeq2 (73). DEGs were identified using the following thresholds: adjusted p-value < 0.5 and absolute log_2_ fold change > 1. The z-scores of gene expressed matrix were used for heatmap plotting. DEGs were sent for GO analysis, which was performed using DAVID (70).

## Data availability

The NCBI GEO accession number for the RNA-seq data reported in this paper is GSE281010.

## Supporting Information

This article contains supporting information, including one figure and three tables.

## Acknowledgments

We are grateful to the Genomics (RRID: SCR_021123) and the Flow Cytometry Cores (RRID: SCR_021134) at Penn State College of Medicine for their support of our research.

## Funding and additional information

This work was partially funded by NIGMS Early-Stage Investigator Maximizing Investigator’s Research Award (ESI MIRA, R35 GM133496) to Z.G., Diversity Supplement to ESI MIRA, and the GlaxoSmithKline Graduate Fellowship to J.C. The content is solely the responsibility of the authors and does not necessarily represent the official views of the National Institutes of Health.

## Footnotes

(AP): Alkaline Phosphatase
(ASD): Autism Spectrum Disorder
(DEG): Differentially expressed genes
(ESC): Embryonic stem cell
(EB): Embryoid body
(GO): Gene Ontology
(GREAT): Genomic regions enrichment of annotations tool
(IP): Immunoprecipitation
(KD): Knockdown
(KO): Knockout
(NE): Nuclear extract
(NPC): Neuronal progenitor cell
(O/N): Overnight
(PcG): Polycomb group
(*POGZ*): *POGO-transposable element with ZNF domain*
(pSMAD): Phosphorylated SMAD
(PRC1): Polycomb Repressive Complex 1
(R-SMAD): Regulatory SMAD
(RT): Room temperature
(siRNAs): Short interfering RNAs

## Notes

### Competing Interest Statement

The authors have declared no competing interest.

