## Supplemental Info for "The zinc-finger protein POGZ associates with Polycomb repressive complex 1 to regulate bone morphogenetic protein signaling during neuronal differentiation"

**Materials Included**

Table S1. Raw data of mass spec analysis listing identified proteins from the RING1B IP in neuronal cells.

Figure S1: Stem cell pluripotency is maintained following *Pogz* KO.

Table S2: PRC1.6-POGZ down-regulated target genes.

Table S3: PRC1.6-POGZ up-regulated target genes.

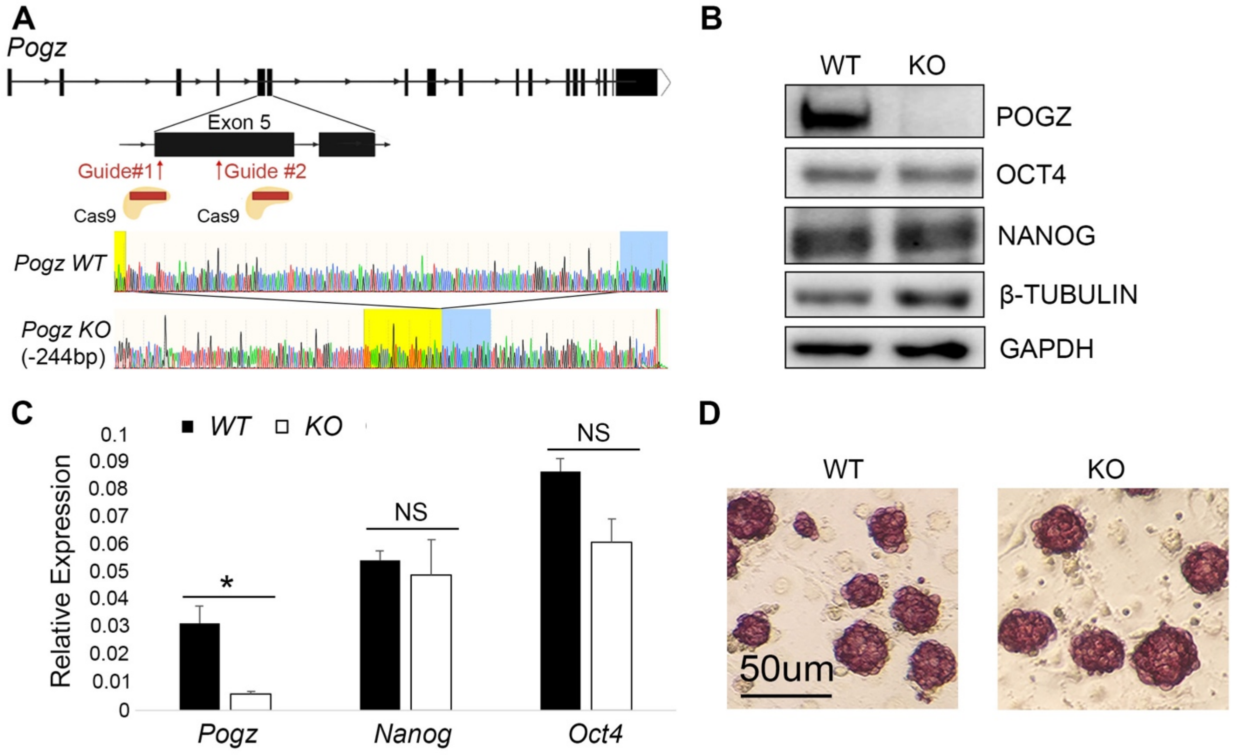


**Figure S1. Stem cell pluripotency is maintained following *Pogz KO***

**A.** (Top) Schematic illustration of the targeting strategy of CRISPR-Cas9 mediated gene editing at the *Pogz* locus to produce *Pogz* KO ESC line. (Bottom) Sanger sequencing results of *Pogz* WT and *Pogz* KO ESC lines displaying the 244 base pair deletion that produced the *Pogz KO* ESC line. **B.** Immunoblotting of *Pogz* WT and KO ESCs to demonstrate *Pogz* KO efficiency and the unaltered protein expression of NANOG and OCT4. **C.** RT-qPCR analysis of *Pogz* and ESC marker genes *Nanog and Oct4* expression in *Pogz* WT and KO ESCs. All mean values and standard deviations were calculated from three independent measurements. Error bars represent standard deviation. * $p<0.05$. **D.** Alkaline phosphatase staining in *Pogz* WT and KO ESCs.

### Supplementary Table 2: PRC1.6-POGZ down-regulated target genes.

| Gene | baseMean | log2FoldChange | lfcSE | pvalue | padj |
|---|---|---|---|---|---|
| Adgrl1 | 1152.196 | -1.21885 | 0.24109 | 1.44E-07 | 0.00000212 |
| Adgrv1 | 556.218 | -3.89346 | 0.299975 | 1.21E-39 | 2.01E-37 |
| Asb4 | 168.838 | -2.17774 | 0.884712 | 0.00111 | 0.006318 |
| Asf1b | 2993.733 | -1.27116 | 0.183824 | 1.48E-12 | 4.48E-11 |
| Atp8a1 | 498.5783 | -1.2518 | 0.504095 | 0.003455 | 0.016455 |
| Bambi | 1862.005 | -1.74113 | 0.267486 | 1.29E-11 | 3.53E-10 |
| Bcl9 | 436.2314 | -1.47175 | 0.46886 | 0.000364 | 0.002417 |
| Boc | 370.6936 | -1.70692 | 1.037182 | 0.009792 | 0.038663 |
| Cdca7 | 3901.457 | -1.20634 | 0.266723 | 0.00000205 | 0.0000236 |
| Cyp46a1 | 72.14976 | -1.89085 | 1.412867 | 0.01011 | 0.039702 |
| Dach1 | 295.9044 | -2.66517 | 0.561158 | 1.73E-07 | 0.0000025 |
| Dusp4 | 220.0973 | -1.57029 | 0.466969 | 0.000151 | 0.001109 |
| Dusp6 | 1078.312 | -1.64407 | 0.301891 | 1.04E-08 | 1.85E-07 |
| E2f3 | 604.6536 | -1.31709 | 0.313113 | 0.00000746 | 0.0000761 |
| Efna5 | 725.3096 | -1.42269 | 0.627466 | 0.00466 | 0.021013 |
| Efnb2 | 736.1427 | -1.35931 | 0.393759 | 0.000142 | 0.001052 |
| Fam181b | 305.2215 | -2.64691 | 0.618049 | 0.00000158 | 0.0000186 |
| Fuz | 232.5152 | -1.64557 | 0.369626 | 0.00000166 | 0.0000194 |
| Gas1 | 684.6893 | -2.36845 | 0.320981 | 1.86E-14 | 6.9E-13 |
| Gtf2i | 2985.962 | -1.23545 | 0.169745 | 1.14E-13 | 3.89E-12 |
| Hmgb2 | 12422.21 | -1.09629 | 0.140901 | 2.7E-15 | 1.09E-13 |
| Hnrnph1 | 4611.673 | -1.00989 | 0.182939 | 1.47E-08 | 2.52E-07 |
| Kat6b | 1512.672 | -1.3361 | 0.17051 | 1.32E-15 | 5.47E-14 |
| Khdrbs2 | 98.09887 | -1.85983 | 0.734378 | 0.00135 | 0.007499 |
| Lca5l | 39.17013 | -2.15613 | 1.395715 | 0.006453 | 0.027523 |
| Lyrm1 | 176.1872 | -1.18844 | 0.540974 | 0.007881 | 0.032344 |
| Lzts1 | 228.4552 | -1.13315 | 0.55594 | 0.012195 | 0.046102 |
| Marcks | 2516.963 | -1.70387 | 0.219473 | 1.65E-15 | 6.78E-14 |
| Mbtd1 | 1566.718 | -1.0148 | 0.295532 | 0.000249 | 0.00173 |
| Mex3b | 410.0009 | -1.13518 | 0.483955 | 0.005872 | 0.025456 |
| Neurog2 | 55.86825 | -2.99221 | 2.329454 | 0.003525 | 0.016718 |
| Nlk | 678.0979 | -1.23751 | 0.295997 | 0.00000925 | 0.0000924 |
| Nr2f2 | 1001.818 | -1.62835 | 0.515617 | 0.000275 | 0.001882 |
| Olig3 | 67.64345 | -5.01249 | 1.302187 | 0.00000468 | 0.0000498 |
| Pabpn1 | 2017.985 | -1.0354 | 0.223695 | 0.00000157 | 0.0000184 |
| Pbx4 | 251.3226 | -1.7447 | 0.542116 | 0.000203 | 0.001441 |
| Pcbp3 | 795.7636 | -1.12116 | 0.465376 | 0.005251 | 0.023184 |

| | | | | | |
|---|---|---|---|---|---|
| Phc2 | 418.3596 | -1.10046 | 0.395767 | 0.002034 | 0.010571 |
| Pim1 | 1076.032 | -1.18931 | 0.290863 | 0.0000148 | 0.00014 |
| Plagl1 | 2004.758 | -1.42363 | 0.346317 | 0.00000971 | 0.0000966 |
| Ppm1k | 207.8264 | -1.36124 | 0.420773 | 0.000309 | 0.002096 |
| Prrc2b | 1579.164 | -1.021 | 0.235824 | 0.00000643 | 0.0000663 |
| Qdpr | 3056.455 | -1.01269 | 0.261924 | 0.0000476 | 0.000398 |
| Qk | 2587.332 | -1.29826 | 0.417383 | 0.000497 | 0.003178 |
| Qtrt2 | 558.5097 | -1.06476 | 0.388997 | 0.002326 | 0.011818 |
| Rab28 | 1200.705 | -1.19344 | 0.216099 | 1.16E-08 | 2.04E-07 |
| Rfx3 | 309.3199 | -1.39845 | 0.543492 | 0.002143 | 0.011035 |
| Rnf44 | 1190.239 | -1.09256 | 0.192388 | 5.34E-09 | 1.01E-07 |
| Sacs | 214.4806 | -1.99509 | 1.098439 | 0.005285 | 0.023304 |
| Sfrp1 | 2978.986 | -3.7863 | 0.222052 | 2.38E-66 | 1.2E-63 |
| Stox2 | 525.8824 | -1.54889 | 0.613513 | 0.002028 | 0.010547 |
| Tnfaip8 | 523.568 | -1.15458 | 0.524018 | 0.008196 | 0.033457 |
| Tox | 93.62585 | -1.76908 | 1.114738 | 0.009537 | 0.037836 |
| Trio | 361.5791 | -1.04568 | 0.457176 | 0.008097 | 0.033091 |
| Ttyh3 | 115.6342 | -1.91581 | 0.781935 | 0.001562 | 0.008506 |
| Wbp1 | 2783.751 | -1.05237 | 0.190839 | 1.58E-08 | 2.68E-07 |
| Wdr44 | 334.5651 | -1.52148 | 0.549407 | 0.001085 | 0.006201 |
| Zfp367 | 375.2026 | -1.48842 | 0.541779 | 0.001203 | 0.006776 |
| Zfp608 | 226.1085 | -2.94301 | 0.731979 | 0.00000388 | 0.000042 |
| Zfp91 | 661.4172 | -1.05 | 0.309763 | 0.000278 | 0.001903 |

### Supplementary Table 3: PRC1.6-POGZ up-regulated target genes.

| Gene | baseMean | log2FoldChange | lfcSE | pvalue | padj |
|---|---|---|---|---|---|
| Abhd2 | 268.808097 | 1.95408902 | 0.63521517 | 0.00026804 | 0.0018438 |
| Adam10 | 1883.18609 | 1.71345825 | 0.22719201 | 9.6814E-15 | 3.7098E-13 |
| Aldh6a1 | 1219.18285 | 1.64020123 | 0.32186127 | 7.0569E-08 | 1.0912E-06 |
| Anxa5 | 5620.27536 | 2.30428235 | 0.19617771 | 1.1049E-32 | 1.3784E-30 |
| Arntl | 742.679881 | 1.85144892 | 0.37607455 | 1.4968E-07 | 2.1959E-06 |
| Asah1 | 8381.91395 | 1.48809523 | 0.19152124 | 2.4717E-15 | 1.0003E-13 |
| Atf7ip | 1809.90862 | 1.01091949 | 0.20854197 | 5.6603E-07 | 7.3429E-06 |
| B4galt1 | 854.565053 | 2.56020667 | 0.25394904 | 7.5363E-25 | 5.8431E-23 |
| Canx | 17067.8345 | 1.01192902 | 0.10645914 | 9.5827E-22 | 6.4112E-20 |
| Cited2 | 247.810373 | 1.84385024 | 0.41113431 | 1.2045E-06 | 1.4493E-05 |
| Cln8 | 1588.71083 | 1.15303309 | 0.31264138 | 8.1685E-05 | 0.00065128 |
| Cobll1 | 296.403001 | 3.29145285 | 0.80066876 | 2.7179E-06 | 3.0719E-05 |
| Cpeb4 | 110.651657 | 2.64371371 | 1.22262935 | 0.00167577 | 0.00901569 |
| Creb3l2 | 1993.84549 | 2.21938591 | 0.57431127 | 1.3072E-05 | 0.00012617 |
| Ddc | 1160.43496 | 1.47750474 | 0.3014482 | 2.3371E-07 | 3.2779E-06 |
| Dnajc1 | 523.126059 | 1.06847661 | 0.44104168 | 0.00564573 | 0.02461219 |
| Eef2kmt | 762.365787 | 1.06152016 | 0.39353126 | 0.00267591 | 0.01328907 |
| Eif2ak3 | 491.895428 | 1.68868957 | 0.75378148 | 0.00356512 | 0.01687054 |
| Esyt2 | 541.013807 | 1.06774655 | 0.27606795 | 4.4668E-05 | 0.0003764 |
| Etv5 | 5462.24749 | 1.44084203 | 0.24784541 | 1.6044E-09 | 3.2986E-08 |
| F3 | 2684.68016 | 3.32790059 | 0.50057157 | 2.9627E-12 | 8.6851E-11 |
| Fam210b | 1114.49999 | 1.55932499 | 0.30881444 | 1.0192E-07 | 1.5276E-06 |
| Fam76a | 749.197582 | 1.08182102 | 0.32203013 | 0.0003138 | 0.00212337 |
| Flrt3 | 3377.00184 | 1.81456715 | 0.40014033 | 8.3881E-07 | 1.0472E-05 |
| Fuca2 | 796.870575 | 1.08434923 | 0.32890456 | 0.00038056 | 0.00251765 |
| Gad2 | 20.6047582 | 2.77947185 | 2.05168503 | 0.00521383 | 0.02303806 |
| Garnl3 | 258.268647 | 1.15482926 | 0.54980208 | 0.01058589 | 0.04120421 |
| Ggct | 348.502922 | 1.59249355 | 0.67359475 | 0.00295262 | 0.01437498 |
| Gm2a | 993.582586 | 1.88646427 | 0.25884045 | 5.1205E-14 | 1.8154E-12 |
| Hspa5 | 30787.2581 | 1.222004 | 0.15347211 | 6.0435E-16 | 2.5844E-14 |
| Ing2 | 448.938637 | 1.12761335 | 0.47392702 | 0.00578552 | 0.02512913 |
| Insig1 | 1997.65345 | 2.15402895 | 0.35621817 | 1.9507E-10 | 4.4952E-09 |
| Kctd9 | 482.311372 | 1.07610981 | 0.42238191 | 0.00399641 | 0.01853745 |
| Klf6 | 1556.80457 | 3.13514039 | 0.26839259 | 1.2539E-32 | 1.5526E-30 |
| Klf7 | 339.649791 | 1.21736023 | 0.41791708 | 0.00111535 | 0.00634633 |
| Myh9 | 1633.38088 | 1.69095959 | 0.25999554 | 1.6331E-11 | 4.3916E-10 |
| Nabp1 | 430.400607 | 2.6020291 | 0.49196821 | 1.2303E-08 | 2.1375E-07 |

| | | | | | |
|---|---|---|---|---|---|
| Nub1 | 1771.36964 | 1.29046065 | 0.27299596 | 7.0867E-07 | 8.9964E-06 |
| Peg10 | 461.036509 | 1.20677542 | 0.45162487 | 0.00221371 | 0.01132587 |
| Plekhf2 | 2579.32435 | 1.01659083 | 0.25435376 | 2.8338E-05 | 0.00025063 |
| Polg | 1824.50295 | 1.59738712 | 0.34349643 | 7.1124E-07 | 9.0099E-06 |
| Por | 1814.57748 | 1.21164877 | 0.21948449 | 1.1964E-08 | 2.0917E-07 |
| Prkab2 | 480.180114 | 2.10764685 | 0.3625846 | 8.7072E-10 | 1.8569E-08 |
| Rab1a | 8039.02092 | 1.38946455 | 0.21251288 | 1.7785E-11 | 4.7519E-10 |
| Ralgapa2 | 507.569449 | 1.42304381 | 0.39505041 | 7.8557E-05 | 0.00062755 |
| Rbm15 | 438.763164 | 1.07128745 | 0.31125059 | 0.00022973 | 0.00161033 |
| Rcan1 | 1698.35824 | 2.0552304 | 0.41972273 | 1.2565E-07 | 1.8631E-06 |
| Rell1 | 499.035956 | 1.42608996 | 0.43641457 | 0.00025862 | 0.00178643 |
| Retreg1 | 2011.03305 | 4.18678919 | 0.20173208 | 9.4194E-97 | 9.1933E-94 |
| Rfk | 2860.26418 | 1.84248282 | 0.19970451 | 5.3093E-21 | 3.4012E-19 |
| Ripk2 | 290.228813 | 1.61272037 | 0.69367775 | 0.00317971 | 0.0153544 |
| Rrbp1 | 2163.72735 | 1.61750779 | 0.36537396 | 2.0054E-06 | 2.3139E-05 |
| Sde2 | 2827.51282 | 1.27648627 | 0.14770001 | 1.8638E-18 | 9.8802E-17 |
| Sec23a | 3890.25081 | 1.28378192 | 0.21154434 | 4.6469E-10 | 1.0212E-08 |
| Sel1l | 2080.01342 | 1.21843752 | 0.35823276 | 0.00021735 | 0.00153136 |
| Shroom3 | 1177.59307 | 1.17733766 | 0.30700935 | 4.4317E-05 | 0.00037382 |
| Slc19a2 | 3188.4801 | 1.1394681 | 0.31501872 | 0.00010895 | 0.00083416 |
| Slc1a4 | 324.398344 | 2.05274019 | 0.54122556 | 1.9476E-05 | 0.00017863 |
| Specc1 | 288.579259 | 2.18190649 | 0.77348878 | 0.0004785 | 0.00307607 |
| Sptlc2 | 2593.05313 | 1.71401165 | 0.43874424 | 1.6105E-05 | 0.00015122 |
| Stat3 | 3415.23247 | 1.07132438 | 0.2134066 | 2.2359E-07 | 3.1546E-06 |
| Stt3a | 12157.5323 | 1.70999198 | 0.22700952 | 1.0516E-14 | 3.9927E-13 |
| Tax1bp3 | 3687.71503 | 1.72950687 | 0.1852144 | 2.0458E-21 | 1.3416E-19 |
| Tpcn1 | 152.824517 | 1.93596049 | 0.80418835 | 0.00179282 | 0.00953414 |
| Uap1 | 1270.19301 | 1.47827023 | 0.35334816 | 6.9057E-06 | 7.0815E-05 |
| Ubald2 | 994.038027 | 1.03389408 | 0.19254611 | 3.5455E-08 | 5.7224E-07 |
| Umad1 | 253.604844 | 1.35890559 | 0.56518021 | 0.00381642 | 0.01789769 |
| Wrnip1 | 973.40727 | 1.12825715 | 0.31615553 | 0.00013163 | 0.00098508 |
